# A VLP-based vaccine targeting ANGPTL3 lowers plasma triglycerides in mice

**DOI:** 10.1101/2021.03.08.434130

**Authors:** Alexandra Fowler, Maureen Sampson, Alan T. Remaley, Bryce Chackerian

**Affiliations:** Department of Molecular Genetics and Microbiology, University of New Mexico School of Medicine, MSC08-4660, Albuquerque, NM 87131, USA; Lipoprotein Metabolism Section, Cardio-Pulmonary Branch, National Heart, Lung and Blood Institute, National Institutes of Health, Building 10-2C433, 10 Center Drive, MSC 1666, Bethesda, MD 20892, USA

**Keywords:** ANGPTL3, Virus-like particles, hypertriglyceridemia

## Abstract

Elevated triglycerides (TGs) are an important risk factor for the development of coronary heart disease (CHD) and in acute pancreatitis. Angiopoietin-like proteins 3 (ANGPTL3) and 4 (ANGPTL4) are critical regulators of TG metabolism that function by inhibiting the activity of lipoprotein lipase (LPL), which is responsible for hydrolyzing triglycerides in lipoproteins into free fatty acids. Interestingly, individuals with loss-of-function mutations in ANGPTL3 and ANGPTL4 have low plasma TG levels, have a reduced risk of CHD, and are otherwise healthy. Consequently, interventions targeting ANGPTL3 and ANGPTL4 have emerged as promising new approaches for reducing elevated TGs. Here, we developed virus-like particle (VLP) based vaccines that target the LPL binding domains of ANGPTL3 and ANGPTL4. ANGPTL3 VLPs and ANGPTL4 VLPs are highly immunogenic in mice and vaccination with ANGPTL3 VLPs, but not ANGPTL4 VLPs, was associated with reduced steady state levels of TGs. Immunization with ANGPTL3 VLPs rapidly cleared circulating TG levels following an oil gavage challenge and enhanced plasma LPL activity. These data indicate that targeting ANGPTL3 by active vaccination is potential alternative to other ANGPTL3-inhibiting therapies.

**Highlights:**

- ANGPTL3 and ANGPTL4 are mediators of lipoprotein metabolism that inhibit lipoprotein lipase (LPL) activity.
- Vaccination using virus-like particles (VLPs) targeting ANGPTL3 and ANGPTL4 elicits high-titer IgG antibody responses.
- Immunization with ANGPTL3 VLPs lowers steady-state plasma triglycerides and enhances LPL activity.

## Introduction

Cardiovascular diseases, which include coronary heart disease (CHD), are the leading cause of death globally [1–3]. CHD is caused by the accumulation of plaque narrowing and hardening the arteries, leading to inefficient blood flow to the heart. Although multiple factors contribute to an increased risk of CHD, dyslipidemia is a leading risk factor. An atherogenic serum lipid profile is characterized by one or more of the following changes: elevated low-density lipoprotein cholesterol (LDL-C), high-density lipoprotein cholesterol (HDL-C), and elevated serum triglycerides (TG). Deposition of cholesterol from LDL into the vessel wall is known to be causally related to atherosclerosis, and interventions to manage elevated LDL-C levels in patients have successfully led to an overall decrease in mean patient LDL-C levels in the United States over the past four decades [4]. However, over the same period, mean levels of serum TGs in Americans have increased and may account for some of the residual risk from dyslipidemias after statin treatment. It is estimated that nearly one-third of the adult US population has hypertriglyceridemia [4]. TGs are primarily involved in energy metabolism and are transported through the bloodstream in the form of circulating lipoproteins, such as chylomicrons, low-density lipoproteins (LDL) and very low-density lipoproteins (VLDL). These lipoproteins can accumulate in the inner lining of the arteries and provoke an inflammatory state in which immune cells are recruited to these sites and further lead to the development of atherosclerosis [5, 6]. Importantly, elevated TGs are an independent factor associated with an increased risk for CHD, irrespective of LDL-C or HDL-C levels [7, 8].

Hypertriglyceridemia can be treated by lifestyle changes (such as modifying diet and/or increasing exercise) or by pharmacotherapy. The most common medications that are prescribed for lowering circulating lipids include statins, which primarily lowers LDL-C and is not that effective for TGs. Other drugs, such as fibrates, niacin, and omega-3 fatty acids can more effectively lower TGs but there is uncertainty if they can reduce cardiovascular events. These therapies are often used in combination; however, even combination therapies are often ineffective at substantially reducing TG levels in patients with moderate to severe hypertriglyceridemia, leaving these individuals at risk of developing CHD [9] and sometimes acute pancreatitis in cases of severe hypertriglyceridemia. This inefficiency can be attributed to multiple factors, including adherence to therapy or statin intolerance, in which patients discontinue medication due to the development of adverse effects and/or myalgia [10, 11], but also reflects the limitation of these existing therapeutics. Hence, additional next-generation lipid-lowering therapies that effectively lower circulating TGs levels are needed.

Circulating triglyceride rich lipoproteins (TRLs) are cleared within the capillaries through an enzymatic reaction involving lipoprotein lipase (LPL), an enzyme that is responsible for hydrolyzing TGs into free fatty acids as an energy source for the surrounding adipocytes or myocytes. Members of the angiopoietin-like (ANGPTL) protein family, including ANGPTL3, ANGPTL4 and ANGPTL8, are key regulators of LPL activity. These proteins bind to LPL and inhibit its activity, slowing TG metabolism and leading to an increase of TGs in circulation [12]. Loss of function (LOF) mutations in the genes that encode ANGPTL3 and ANGPTL4 are associated with hypertriglyceridemia in mice [13–15] and familial combined hypolipidemia in humans [16, 17], which is characterized by low levels of LDL, HDL, and TGs, and protection against CHD [18, 19].

The association between loss-of-function mutations in human ANGPTL3 and ANGPTL4 and a reduced risk of CHD has suggested that these proteins may be promising therapeutic targets for lowering circulating TGs. Monoclonal antibodies (mAbs) against both ANGPTL3 [20] and ANGPTL4 [14] have been shown to reduce circulating levels of TGs in mice, and, clinical trials of Evinacumab, a fully humanized mAb against ANGPTL3, have shown that this drug can significantly reduce circulating TGs and LDL-C levels in patients with familial hypercholesterolemia [19, 21–23]. Although these data are encouraging, the use of mAbs to treat chronic disease has some important shortcomings that may limit patient access and affect clinical efficacy. These include the need for frequent injections of high doses of antibody, the expense associated with mAb-based therapies, and the possibility of inducing anti-drug antibodies, a phenomenon that occurs even with fully human mAbs. As an alternative approach, we assessed whether an active vaccination approach for targeting ANGPTL molecules could be an effective strategy for lowering TG levels.

We and others have previously shown that virus-like particle (VLP)-based vaccines can be used as platform technologies to generate strong antibody responses against self-proteins [24–27]. Although the mechanisms of immunological tolerance usually limit the induction of antibodies against self, B-cell tolerance can be overcome by using vaccines which display self-antigens in the high repetitive, multivalent structural context of a VLP [24, 28]. These data raised the possibility that vaccination against self-molecules involved in disease could be an effective, long-lasting, and economical alternative to mAb therapies [29]. Previously, we showed that a VLP-based vaccine targeting the self-antigen proprotein convertase subtilisin/kexin type 9 (PCSK9), a molecule involved in cholesterol homeostasis, could elicit high-titer anti-PCSK9 antibodies in mice and non-human primates and reduce circulating cholesterol levels [30]. Here, we investigated the ability of bacteriophage VLP-based vaccines to target the LPL-inhibiting domains of ANGPTL3 and ANGPTL4. We show that vaccination with Qβ bacteriophage VLPs displaying an ANGPTL3 peptide are highly immunogenic, lower TG levels and increase LPL activity in mice.

## 2. Methods and materials

### 2.1. Vaccines

Qβ bacteriophage VLPs were produced in *Escherichia coli* (*E.coli*) using methods previously described for the production of other RNA bacteriophages [31]. A peptide representing the mouse ANGPTL3 amino acids 32-47 (^32^EPKSRFAMLDDVKILA^47^), modified to include a C-terminal cysteine residue preceded by a 3-glycine-spacer sequence (-GGGC), was synthesized by GenScript (Piscataway, NJ). A peptide representing the mouse ANGPTL4 amino acids 29-53 (^29^QPEPPRFASWDEMNLLAHGLLQLGH^53^) was also synthesized by Genscript) with the same C-terminal-GGGC modification. The ANGPTL3 and ANGPTL4 peptides were conjugated to Qβ bacteriophage VLPs using succinimidyl 6-((beta-maleimidopropionamido)hexanoate), SMPH; Thermo Fisher Scientific, Waltham, MA), as described previously [32]. The efficiency of peptide-VLP conjugation was measured using sodium dodecyl sulfate-polyacrylamide gel electrophoresis (SDS-PAGE).

### 2.2. Animals and immunizations

All animal studies were performed in accordance with guidelines of the University of New Mexico Animal Care and Use Committees (protocol 19-200870-HSC). Mouse immunization experiments were performed using 4-week-old Balb/c mice obtained from The Jackson Laboratory (Bar Harbor, ME). Mice were immunized three times at 3-week intervals with 5μg doses of ANGPTL3_32-47_-Qβ VLPs, ANGPTL4_29-53_-Qβ VLPs, or wild-type Qβ VLPs, or a mixture of ANGPTL3_32-47_-Qβ VLPs and ANGPTL4_29-53_-Qβ VLPs (5μg of each vaccine). Blood plasma was collected via retroorbital bleed prior to the first immunization and two weeks following each immunization. A final cardiac bleed was performed three weeks following the third immunization. Mice were fasted for 3 hours prior to each bleed.

### 2.3. Measuring antibodies

ANGPTL3 and ANGPTL4-specific IgG were detected by direct ELISA, using either ANGPTL3 or ANGPTL4 peptides (representing the mouse sequences), recombinant proteins representing either full-length mouse ANGPTL3 or the N-terminal domain of mouse ANGPTL3 (R&D Systems, Minneapolis, MN), or the N-terminal domain of human ANGPTL4 (R&D Systems) as the target antigens. ELISAs were performed as described [30] with a few modifications. Sera was diluted serially, using 4-fold dilutions starting at 1:40. The wells were probed with horseradish peroxidase (HRP)-conjugated secondary antibody (goat anti-mouse-IgG [Jackson ImmunoResearch, West Grove, PA; diluted 1:4000]) for 1 hour. The reaction was developed using TMB (Thermo Fisher Scientific) and stopped using 1% HCl. The optical density was measured at 450nm (OD450) using an accuSkan FC microwell plate reader (Thermo Fisher Scientific).

### 2.4. Plasma lipid quantification

Plasma lipids were measured enzymatically using a ChemWell instrument and Roche reagents.

### 2.5. Oil oral gavage challenge in mice

Olive oil gavages were performed on vaccinated mice three weeks following the third immunization using a protocol modified from [33]. 24 hours prior to gavage, blood plasma was collected from mice in order to monitor baseline TG levels. For 12 hours prior to gavage and over the course of the procedure mice were fasted. At gavage, each mouse received 300μL of olive oil by oral gavage. Briefly, a straight feeding needle (18 gauge x 2 inches) was pre-filled with olive oil, mice were then held in an upright position, the needle was placed in the back of the mouth in order to reach the esophagus, and the oil was slowly administered. Blood plasma was collected 3 hours post-gavage by retroorbital bleed and 6 hours post-gavage by terminal cardiac bleed.

### 2.6 Measuring LPL activity

LPL levels were quantitated in the plasma of vaccinated mice. In order to release LPL into the blood, mice were treated with 70 Units of heparin (diluted in PBS to a total volume of 150μL) by tail vein injection. Ten minutes following injection, blood plasma was collected by retroorbital bleed. LPL activity was measured by using an LPL Activity Assay Kit (Abcam, Cambridge, MA), following the manufacturer’s instructions. Fluorescence was measured at 482nm (excitation)/515nm (emission) on a microplate reader (HTX Microplate Reader; Biotek, Winooski, VT) in kinetic mode, measuring every 10 minutes for 1 hour at 37°C. LPL activity was quantitated by comparing experimental samples (tested in duplicate using 8μL of blood plasma) to an internal standard curve.

### 2.7 Statistical Analysis

All statistical analysis of data was performed using GraphPad Prism 9.

## 3. Results

### 3.1. Engineering and Characterization of VLPs displaying epitopes from ANGPTL3 and ANGPTL4

ANGPTL3 and ANGPTL4 have related structural features that are characteristic of other members of the angiopoietin and ANGPTL families. Both proteins possess an N-terminal domain, that includes a region that is predicted to be intrinsically disordered followed by a coiled-coil domain, and a C-terminal fibrinogen-like domain [34]. Using mAbs that inhibit LPL binding, the LPL-inhibiting domains of ANGPTL3 and ANGPTL4 were mapped to the disordered region of the N-terminal domain [35, 36]. We hypothesized that antibodies directed against this region could interfere with the LPL inhibitory activity of ANGPTL3 and ANGPTL4. In an initial pilot study, we engineered VLP-based vaccines that targeted peptides representing the entire LPL-inhibiting domains of ANGPTL3 and ANGPTL4 (Supplemental Fig. 1). Peptides corresponding to ANGPTL3 amino acids 32-55 (ANGPTL3_32-55_) and ANGPTL4 amino acids 29-53 (ANGPTL4_29-53_) were synthesized and chemically conjugated to Qβ bacteriophage VLPs using a chemical crosslinker, SMPH. We were successfully able to conjugate ANGPTL4_29-53_ to VLPs, but the peptide representing ANGPTL3_32-55_ formed insoluble aggregates upon reaction with VLPs. As an alternative approach, we synthesized a shorter peptide representing ANGPTL3 amino acids 32-47 (ANGPTL3_32-47_) and were able to successfully conjugate this peptide to Qβ VLPs. The efficiency of the chemical modification was determined by SDS-PAGE. Based on the efficiency of conjugation to the individual coat protein subunits of the VLPs, we estimate that approximately 270-360 ANGPTL3_32-47_ or ANGPTL4_32-54_ peptides were conjugated to each individual Qβ VLP (Supplemental Fig. 2).

### 3.2. VLPs displaying ANGPTL3 and ANGPTL4 peptides are immunogenic in mice

In order to assess the immunogenicity of conjugated VLPs, groups of male Balb/c mice were immunized with 5μg of ANGPTL3-peptide displaying VLPs (ANGPTL3_32-47_-Qβ VLPs; hereforth referred to as ANGPTL3 VLPs), 5μg of ANGPTL4-peptide displaying VLPs (ANGPTL4_32-54_-Qβ VLPs; ANGPTL4 VLPs), or, as negative control, 5μg of unmodified Qβ VLPs. Mice were immunized three times at 3-week intervals. Antibody responses were initially evaluated by peptide ELISA, using synthetic peptides representing ANGPTL3_32-47_ or ANGPTL4_32-54_ as target antigens. As shown in Fig. 1, mice immunized with ANGPTL3 VLPs (Fig. 1A) and ANGPTL4 VLPs (Fig. 1B) elicited high titer IgG antibodies against the cognate peptides, as compared to control groups.

**Fig. 1.**
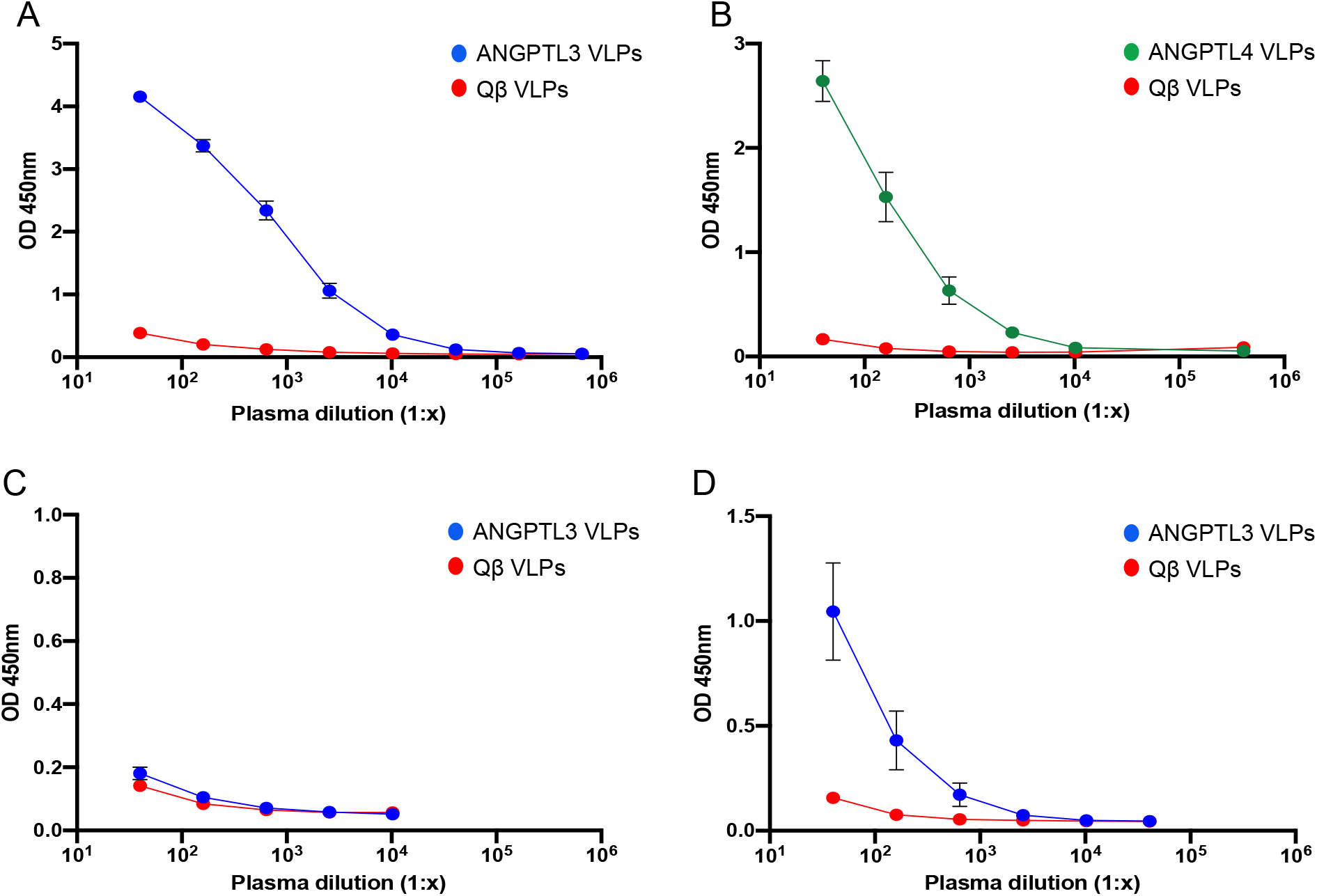
Antibody responses induced by VLP-based vaccines targeting ANGPTL3 and ANGPTL4. Groups of mice (n=8) were immunized with ANGPTL3 VLPs (blue symbols), ANGPTL4 VLPs (green symbols), or unmodified Qβ VLPs (red symbols). Sera was collected from mice three weeks following their third immunization and IgG responses against (A) ANGPTL3 peptide, (B) ANGPTL4 peptide, (C) full-length mouse ANGPTL3, and (D) the N-terminal region of mouse ANGPTL3 were measured by ELISA. Mean OD450 values for each group of mice are shown, error bars represent standard error of the mean (SEM).

ANGPTLs are proteolytically cleaved by proprotein convertases to produce two fragments representing the N-terminal domain and the C-terminal fibrinogen-like domain. The cleaved N-terminal fragments of ANGPTL3 and ANGPTL4 are found in circulation and has enhanced LPL-inhibiting activity compared to full length protein [37, 38]. To assess the potential for anti-ANGPTL3 antibodies elicited following immunization with ANGPTL3 VLPs to bind these active forms of ANGPTL3, we performed ELISAs using either full length ANGPTL3 or the N-terminal fragment (ANGPTL3_17-220_) as target antigens. As shown in Fig. 1C, sera from mice immunized with ANGPTL3 VLPs failed to bind to fulllength mouse ANGPTL3 protein. However, these antibodies bound strongly to mouse ANGPTL3_17-220_ (Fig. 1D), suggesting that they may have the ability to functionally neutralize the more enzymatically active form of ANGPTL3.

### 3.4. Immunization with ANGPTL3 VLPs, but not ANGPTL4 VLPs, lower resting triglyceride levels in mice

In order to test the functional effects of immunization with ANGPTL3 or ANGPTL4 Qβ VLPs, we measured lipid levels in the plasma of vaccinated mice prior to the initial vaccination and three weeks following the third immunization. Control mice immunized with wild-type Qβ VLPs showed an increase in TG levels over the course of the study, which likely reflect natural increases in TG levels as the mice age. In contrast, TG levels in mice immunized with ANGPTL3 VLPs were reduced at the post-immunization timepoint, 34% lower than mice immunized with control VLPs (Fig. 2A). In contrast, immunization with ANGPTL4 VLPs did not reduce TG levels relative to controls (Supplemental Fig. 3). We also assessed total cholesterol levels and found that mice immunized with ANGPTL3 VLPs had similar levels as mice immunized with control VLPs (Fig. 2B), which perhaps reflects the fact that mice carry most of their cholesterol on HDL. Thus, these data indicate that immunization with ANGPTL3 VLPs, but not ANGPTL4 VLPs, lowers steady-state TG levels in mice.

**Fig. 2.**
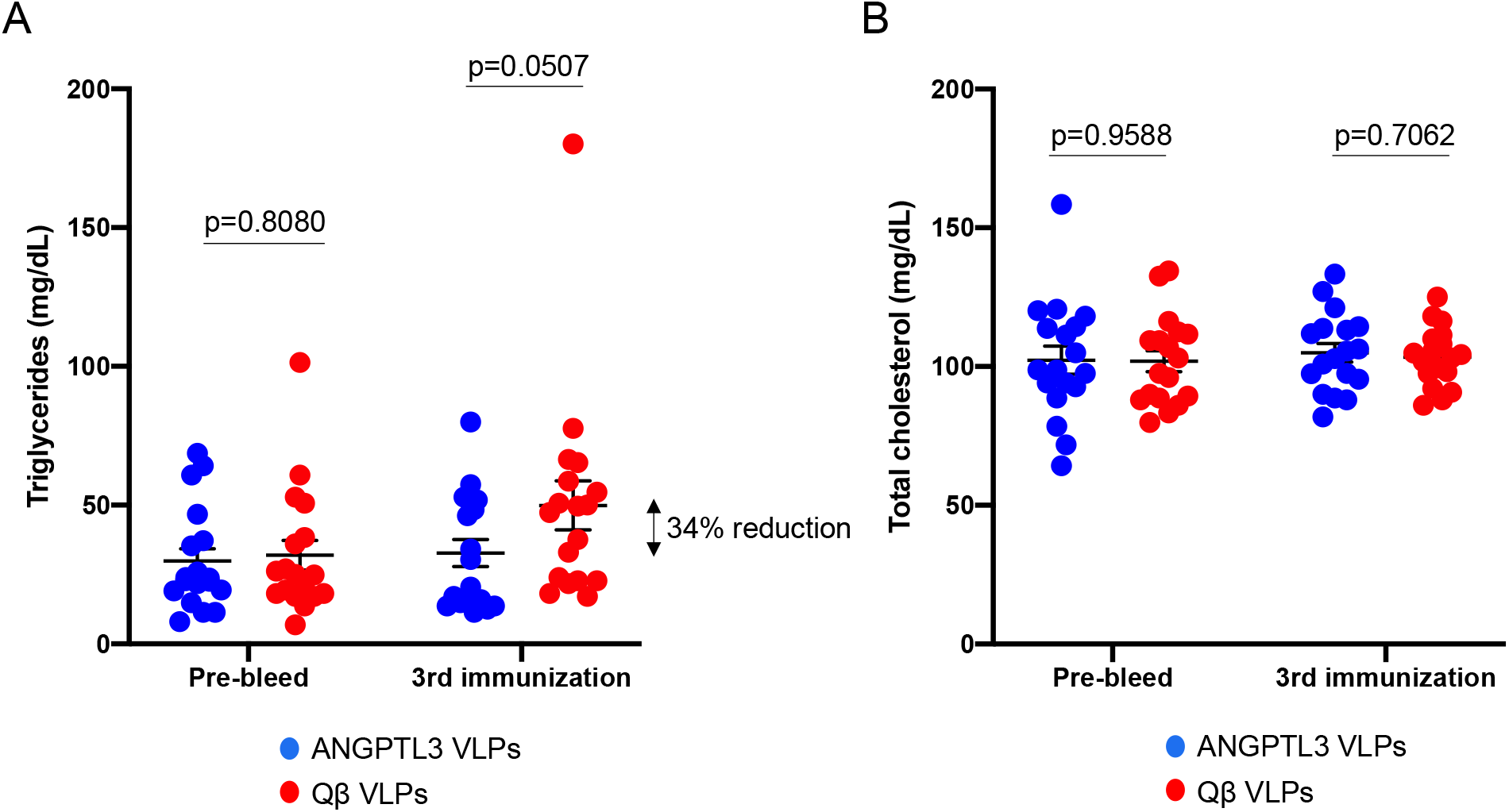
Triglyceride (TG) and total cholesterol levels in mice vaccinated with ANGPTL3 VLPs. Groups of mice (n=18, from two independent experiments) were immunized three times with ANGPTL3 VLPs (blue symbols) or unmodified Qβ VLPs (red symbols). Plasma (A) TG and (B) total cholesterol levels were measured prior to immunization and three weeks following the third immunization. Each data point represents an individual mouse, means and SEM are also represented on each graph. Lipid levels were compared statistically by unpaired t-test.

### 3.5. Mice immunized with ANGPTL3 Qβ VLPs are able to rapidly hydrolyze triglycerides upon lipid challenge

As another measure to assess ANGPTL3/ANGPTL4 inhibition, we evaluated the ability of vaccinated or control mice to clear bolus TRLs introduced by oral gavage. Three weeks after their third vaccination, fasted mice were administered olive oil by gavage in order to induce a postprandial hypertriglyceridemia (PHTG) state [39]. Plasma was collected 24 hours prior to gavage (in order to assess steady state TG levels), three hours post-gavage (to monitor peak TG levels), and six hours post-gavage (to measure the rate of TG clearance in the different groups). As shown in Fig. 3A, mice immunized with ANGPTL3 VLPs had lower steady-state TG levels than control mice at the pre-gavage timepoint, similar to what was observed in Fig. 2A. Both ANGPTL3 VLP- and control-immunized mice had similar peak TG levels at three hours post-gavage. However, mice immunized with ANGPTL3 VLPs cleared TGs more rapidly than control mice; we measured a 47% reduction of TG levels at 6 hours post-gavage in the ANGPTL3 VLP-immunized group relative to controls. In contrast, mice immunized with ANGPTL4 VLPs did not hydrolyze TGs more rapidly than control mice (Fig. 3B).

**Fig. 3.**
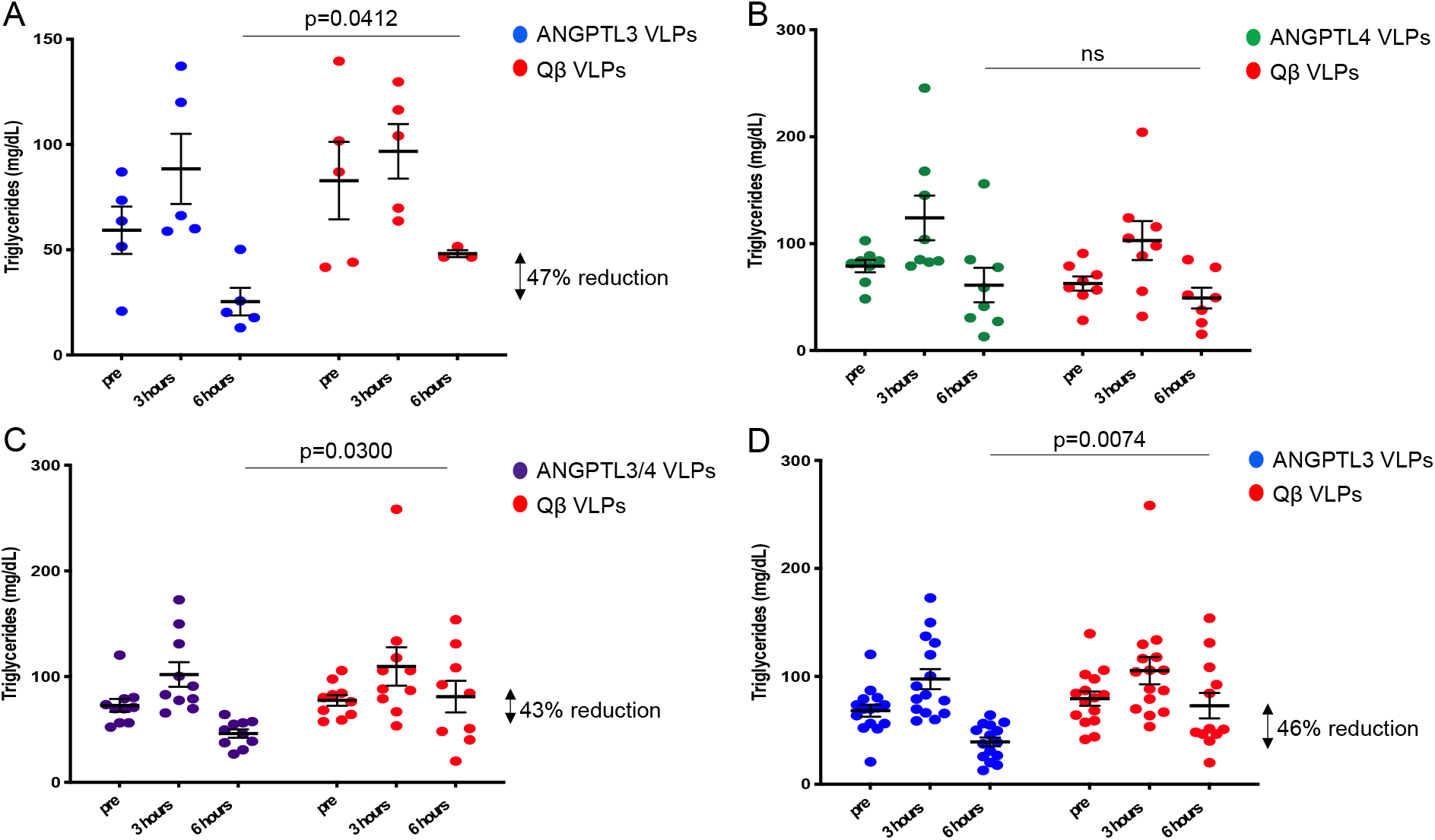
Triglyceride (TG) levels in immunized mice that received an olive oil gavage. Groups of mice were immunized three times with (A) ANGPTL3 VLPs (n=5; blue symbols), (B) ANGPTL4 VLPs (n=8; green symbols), or (C) a mixture of ANGPTL3 and ANGPTL4 VLPs (n=10; purple symbols), or control, unmodified Qβ VLPs (red symbols). Three weeks following their final immunization, mice were given an olive oil gavage. Plasma was collected 24 hours prior to gavage, 3 and 6 hours post-gavage, and then TG levels were measured. Each data point represents an individual mouse, and means and SEM are also represented on each graph. Lipid levels at the 6 hours post-gavage timepoint were statistically compared by unpaired t-test; ns denotes not significant. Panel (D) shows aggregated data from all mice immunized with ANGPTL3 VLPs compared to all mice immunized with control Qβ VLPs (n=15 per group).

Although ANGPTL3 and ANGPTL4 both play roles in lipid metabolism through their LPL-inhibiting activity, they do differ by the nutritional state at which they are expressed. While ANGPTL3 is unaffected by the nutritional environment and expression is induced during either fasted or fed states, ANGPTL4 is primarily expressed during the fasted state [40]. We hypothesized that blocking both ANGPTL3 and ANGPTL4 may work synergistically to rapidly lower TGs in mice following oil oral gavage challenge. To test this hypothesis, a group of male Balb/c male mice were immunized three times with a mixture of ANGPTL3 VLPs (5μg) and ANGPTL4 VLPs (5μg). This combination vaccine induced high anti-ANGPTL3 and anti-ANGPTL4 VLPs peptide titers, similar to the antibody levels induced by each vaccine individually (data not shown). Upon olive oil oral gavage, mice immunized with the combination ANGPTL3/4 VLP vaccine had a significant reduction in TG levels at the 6-hour time point compared to the mice immunized with Qβ VLPs (Fig. 3C). However, this reduction was similar to that observed in mice immunized with ANGPTL3 VLPs alone, suggesting that antibodies against ANGPTL3 accounted for this effect. To more fully assess the effectiveness of immunization with ANGPTL3 VLPs on TG metabolism, data from all mice immunized with ANGPTL3 VLPs was aggregated to and then depicted in Fig. 3D. Taken together, these data indicate that mice immunized with ANGPTL3 VLPs are able to efficiently and rapidly clear the high levels of circulating TGs in an induced postprandial state.

### 3.6. Immunization with ANGPTL3 VLPs increases plasma LPL activity

Humans who are homozygous for a complete ANGPTL3 deficiency have higher plasma LPL activity than heterozygotes and non-carriers [41], suggesting that this may also be true for ANGPTL3-vaccinated mice. Mice were immunized with ANGPTL3 VLPs, Qβ VLPs, or as an additional negative control, mock-immunized with PBS. Following three immunizations, mice were treated with heparin, which liberates LPL from the luminal surface of vascular endothelial cells [42], and then plasma was collected. LPL activity was measured using an *in vitro* assay which measures the amount of a quenched substrate that is hydrolyzed by LPL. As shown in Fig. 4, LPL activity was significantly increased in the mice immunized with ANGPTL3 VLPs compared to the control groups. These data indicate that immunization with ANGPTL3 VLPs elicits anti-ANGPTL3 antibodies that bind to the LPL-inhibiting domain of ANGPTL3, resulting in an increase in LPL activity in the plasma of mice.

**Fig. 4.**
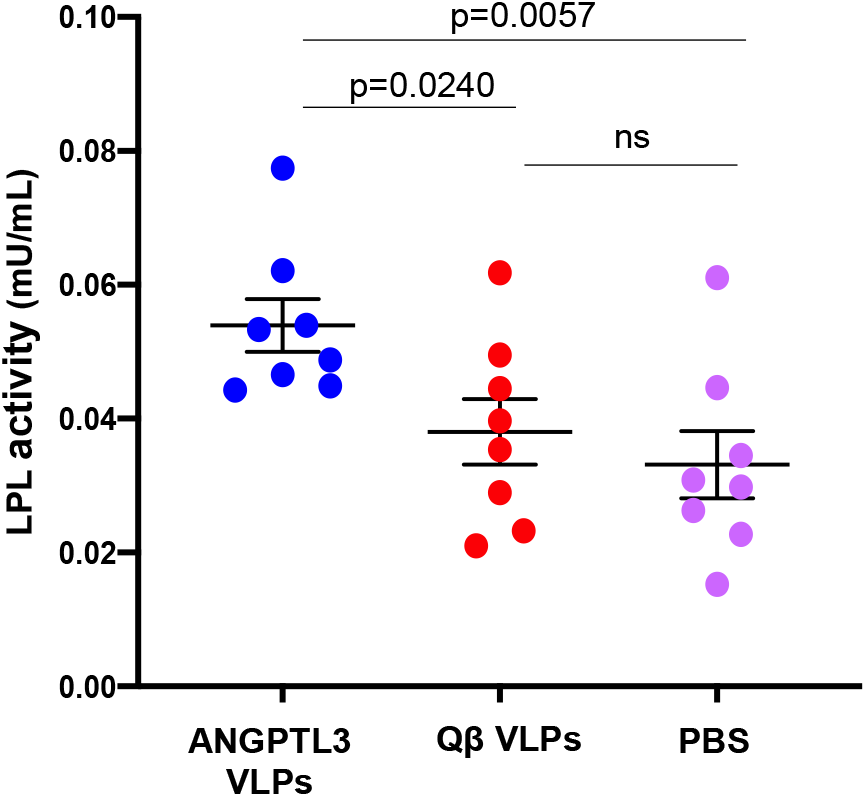
Enhancement of LPL activity in mice vaccinated with ANGPTL3 VLPs. Groups of mice (n=8) were immunized three times with ANGPTL3 VLPs (blue symbols), unmodified Qβ VLPs (red symbols), or PBS (pink symbols). Three weeks following the final immunization, mice received a tail-vein injection of heparin (~70 Units), plasma was collected 10 minutes later, and then LPL activity from individual plasma samples was measured using an *in vitro* assay. One Unit of LPL activity is equal to the amount of LPL that generates 1.0 nmol of fatty acid product per minute (at pH 7.4 and 37°C). Samples were tested in duplicate and then averaged. Groups were compared using a two-tailed, unpaired t-test.

## 4. Discussion and Conclusion

Over the past 20 years, genetic studies have uncovered a number of secreted proteins that play key roles in regulating circulating lipid levels. These include PCSK9, which regulates expression of the low-density lipoprotein receptor (LDL-R) and is a negative regulator of LDL-C homeostasis. Another class of regulatory secretory proteins are members of the ANGPTL family, including ANGPTL3 and ANGPTL4, which control the metabolism of TGs by inhibiting LPL activity. Loss of function mutations in PCSK9, ANGPTL3, and ANGPTL4 are associated with reduced LDL-C and TG levels, and also a reduced risk for developing CHD [18, 43–45]. Given their critical roles in controlling the levels of proatherogenic lipids, and the remarkable fact that individuals with inactivating mutations in PCSK9, ANGPTL3 and ANGPTL4 are otherwise healthy, these molecules have emerged as exciting therapeutic targets for reducing the risk of CHD. PCSK9 inhibitors, which include the mAbs evolocumab and alirocumab, were first approved for use in the United States in 2015 and have been shown to dramatically reduce LDL-C levels in patients with hypercholesterolemia. More recent studies have focused on the development of ANGPTL3 inhibitors. These include Evinacumab, a fully humanized mAb targeting ANGPTL3, and vupanorsen, an antisense oligonucleotide (ASO) that targets hepatic ANGPTL3 mRNA for degradation. Evinacumab has been shown to reduce plasma lipids in animal models [46] and, subsequently, in human clinical trials [19, 21–23]. A phase 3 trial in patients with homozygous familial hypercholesterolemia showed that treatment with Evinacumab led to a 47.1% reduction in LDL levels compared to the placebo control patients [23]. Vupanorsen (also called ANGPTL3-L_Rx_) was developed by Ionis Pharmaceuticals and consists of a GalNAc-conjugated ASO that targets human ANGPTL3 mRNA. In phase 1 and 2 clinical trials, individuals treated with vupanorsen exhibited reduced levels of TGs, LDL, VLDL, as well as decreased *Angptl3* mRNA expression compared to the placebo control group [47, 48]. Collectively, these data suggest that decreasing the function or expression of ANGPTL3 can result in decreased lipid levels.

Here, we employed a different strategy by developing an epitope-targeted VLP-based vaccine that induces antibodies that bind to a region of ANGPTL3 that is known to be involved in the inhibition of LPL activity. This vaccine circumvents the mechanisms of B cell tolerance by displaying the ANGPTL3 epitope multivalently on a VLP surface and by linking the targeted self-peptide to foreign T helper epitopes provided by the VLP platform. We demonstrated that immunization with ANGPTL3 VLPs induced high-titer antibodies that bound to the functionally active N-terminal fragment of ANGPTL3, and that these antibodies lowered steady-state TG levels in mice. Moreover, vaccination with ANGPTL3 VLPs led to increased LPL activity *in vivo* and more rapid hydrolysis of TGs upon olive oil gavage. Since ANGPTL3 inhibits the catalytic activity of LPL, these findings could be explained by the induction of anti-ANGPTL3 antibodies that interfere with the LPL binding domain within ANGPTL3, therefore permitting LPL to function. Taken together, these data suggest that vaccination against ANGPTL3 is a potential alternative to other ANGPTL3-lowering strategies.

We also examined the efficacy of an analogous vaccine targeting the LPL-inhibitory domain of ANGPTL4, but vaccination with ANGPTL4 VLPs had no effect on steady-state TG levels or on TG hydrolysis upon olive oil challenge. This may be due to the fact that while ANGPTL3 in constitutively expressed, ANGPTL4 is primarily expressed during a fasted state. It is also possible that the ANGPTL4 epitope that we targeted is not exposed on the native or cleaved mouse ANGPTL4. Additionally, while the primary function of ANGPTL3 is to bind and inhibit LPL activity, ANGPTL4 is multifunctional, not only playing a role in lipid metabolism, but it also plays functional roles in angiogenesis, inflammation, and vascular permeability [49].

While a vaccine targeting a self-antigen may seem like an unconventional approach, VLP-based vaccines have been previously used to induce high-titer and long-lasting antibody responses against self-antigens involved in chronic disease in numerous preclinical animal models [50–53]. Moreover, several VLP-based vaccines targeting self-antigens have entered human clinical trials and have been shown to be immunogenic and safe [54–57]. An active vaccination approach may have advantage over other, more conventional, drug-based therapies (such as mAbs). Treatment of chronic diseases can often span decades, which places a burden on patients to comply with dosing schedules that usually require regular, in-person treatment. In clinical trials of Evinacumab, for example, patients were administered the mAb intravenously every 4 weeks [23]. In contrast, vaccines will likely require infrequent dosing, which potentially will increase patient compliance. Moreover, while vaccines are usually inexpensive, mAb-based drugs are amongst the most expensive drugs on the market. For example, even in the developed world, health insurance companies have limited access to mAb-based PCSK9 inhibitors, which resulted in an increased risk in cardiovascular events in untreated cohorts [58]. Thus, an effective vaccine-based approach could dramatically enhance the affordability and accessibility, particularly to low-resource parts of the world.

In summary, we investigated the potential of targeting ANGPTL3 through active vaccination using an engineered VLP-based vaccine in mice. The data presented in this paper provides evidence that targeting the self-protein ANGPTL3 using VLPs can induce strong antibody responses that lower TGs in mice. This approach potentially represents a more accessible and affordable alternative to other therapies that are currently in development. Furthermore, if active vaccination against ANGPTL3 shows to be successful in a clinical setting, this will have implications for reducing the global burden of CHD.

## Abbreviations

ANGPTL: Angiopoietin-like protein
LPL: lipoprotein lipase
VLP: Virus-like particle
CHD: coronary heart disease
IgG: Immunoglobulin G
LDL-C: low density lipoprotein cholesterol
HDL-C: high density lipoprotein cholesterol
TG: triglyceride
mAb: monoclonal antibody
ASO: antisense oligonucleotide
PHTG1: postprandial hypertriglyceridemia

## Acknowledgments

This research was funded by NIH grant R01HL131696 and by the Intramural Research Program of NHLBI.

## Conflict of Interest Statement

B.C. has equity stakes in Agilvax, Inc. and Flagship Labs (FL) 72. The other authors have no known competing financial interests or personal relationships that could have appeared to influence the work reported in this paper.

