## Supplemental Figures for "A VLP-based vaccine targeting ANGPTL3 lowers plasma triglycerides in mice"

Mouse ANGPTL3 MHTIKLFLFVVPLVIASRVDPDLSSFDSAPS**EPKS-RFAMLDDVKILANGLLQLGH**GLKD

Mouse ANGPTL4 --MRCAPTAGAA..-LCAATAG.L.AQGR.A**QPEPP...SW.EMNL..H.......**..RE

**Supplemental Fig. 1.** An alignment of the N-terminal regions of mouse ANGPTL3 (accession NP_038941.1) and mouse ANGPTL4 (accession EDL10208.1). Periods denote identity between the two proteins and dashes represent gaps. The colored regions of each protein denote the LPL-inhibitory domains of ANGPTL3 and ANGPTL4, the underlined regions indicate the peptides that synthesized and were conjugated to VLPs.

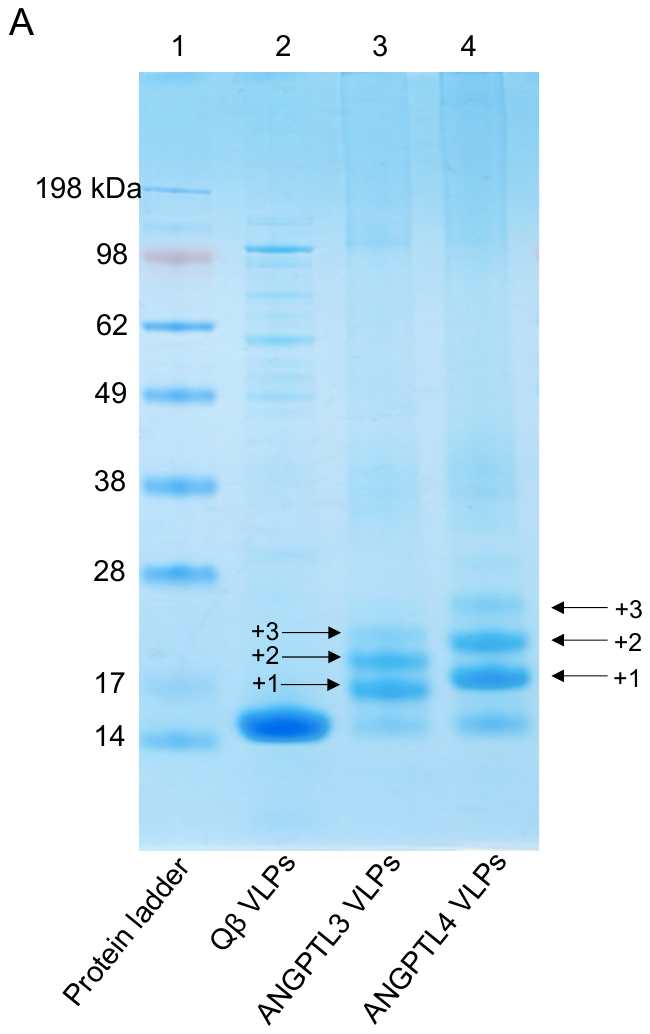

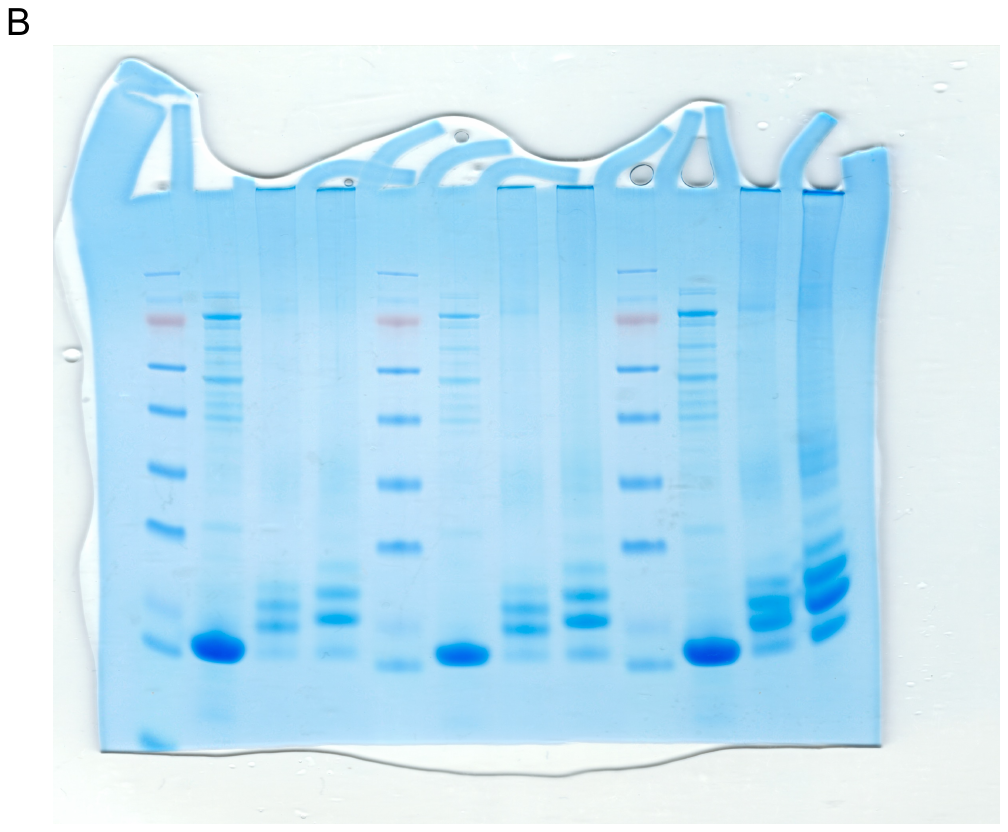

**Supplemental Fig. 2.** (A) SDS-PAGE analysis of unconjugated Qß VLPs (lane 2), ANGPTL3 VLPs (lane 3), and ANGPTL4 VLPs (lane 4). The ladder of bands in the ANGPTL VLP lanes reflect individual copies of coat protein modified with 1, 2, or 3 copies of peptide. Size markers are shown in lane 1. (B) The unmodified gel from which panel A was derived.

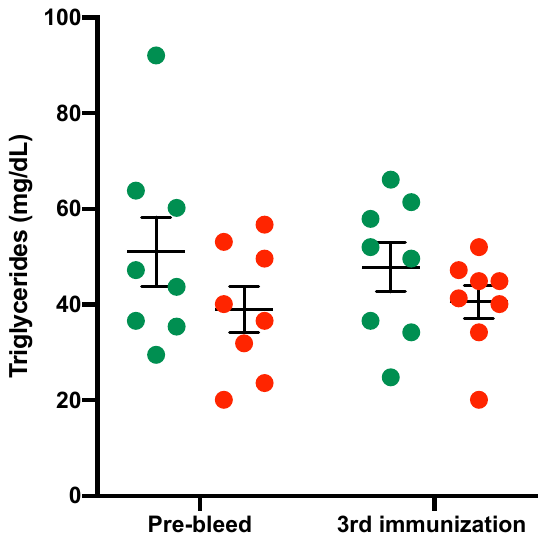

ns

ANGPTL4 VLPs

Qβ VLPs

**Supplemental Fig. 3.** Triglyceride (TG) levels in mice vaccinated with ANGPTL4 VLPs. Groups of mice (n=8, from two independent experiments) were immunized three times with ANGPTL4 VLPs (green symbols) or unmodified Qß VLPs (red symbols). Plasma TG levels were measured prior to immunization and three weeks following the third immunization. Each data point represents an individual mouse, means and SEM are also represented on each graph.
